# Delineation of the First Human Mendelian Disorder of the DNA Demethylation Machinery: *TET3* Deficiency

**DOI:** 10.1101/719047

**Authors:** David B. Beck, Ana Petracovici, Chongsheng He, Hannah W. Moore, Raymond J. Louie, Muhammad Ansar, Sofia Douzgou, Sivagamy Sithambaram, Trudie Cottrell, Regie Lyn P. Santos-Cortez, Eloise J. Prijoles, Renee Bend, Boris Keren, Cyril Mignot, Marie-Christine Nougues, Katrin Õunap, Tiia Reimand, Sander Pajusalu, Muhammad Zahid, Muhammad Arif Nadeem Saqib, Julien Buratti, Eleanor G Seaby, Kirsty McWalter, Aida Telegrafi, Dustin Baldridge, Marwan Shinawi, Suzanne M. Leal, G. Bradley Schaefer, Roger E. Stevenson, Siddharth Banka, Roberto Bonasio, Jill A. Fahrner

**Affiliations:** National Human Genome Research Institute, National Institutes of Health, Bethesda, MD, 20892, USA; Graduate Group in Genetics and Epigenetics, University of Pennsylvania Perelman School of Medicine, Philadelphia, PA, 19104, USA; Department of Cell and Developmental Biology, University of Pennsylvania Perelman School of Medicine, Philadelphia, PA, 19104, USA; Epigenetics Institute, University of Pennsylvania Perelman School of Medicine, Philadelphia, PA, 19104, USA; Hunan Key Laboratory of Plant Functional Genomics and Developmental Regulation, Hunan University, Changsha, 410082 Hunan, P.R. China; Greenwood Genetic Center, Greenwood, SC, 29646, USA; Department of Biochemistry, Faculty of Biological Sciences, Quaid-I-Azam University, 45320, Islamabad, Pakistan; Division of Evolution & Genomic Sciences, School of Biological Sciences, Faculty of Biology, Medicine and Health, University of Manchester, Manchester, M13 9PL, UK; Manchester Centre for Genomic Medicine, St Mary’s Hospital, Manchester University NHS Foundation Trust, Health Innovation Manchester, Manchester, M13 9WL, UK; Department of Otolaryngology, University of Colorado School of Medicine, Aurora, CO, 80045, USA; Assistance Publique-Hôpitaux de Paris, Groupe Hospitalier Pitié-Salpêtrière, Département de Génétique, Paris, 75013, France; Centre de Référence Déficiences Intellectuelles de Causes Rares, Paris, France; Assistance Publique-Hôpitaux de Paris, Armand Trousseau Hospital, Department of Neuropediatrics, Paris, 75012, France; Department of Clinical Genetics, United Laboratories, Tartu University Hospital, Tartu, 50406, Estonia; Department of Clinical Genetics, Institute of Clinical Medicine, University of Tartu, Tartu, Estonia; Chair of Human Genetics, Institute of Biomedicine and Translational Medicine, University of Tartu, Tartu, Estonia; Yale University School of Medicine, Department of Genetics, New Haven, CT, USA; Pakistan Health Research Council, 45320, Islamabad, Pakistan; Program in Medical and Population Genetics, Broad Institute of MIT and Harvard, Cambridge, Massachusetts 02142, USA; Analytic and Translational Genetics Unit, Massachusetts General Hospital, Boston, Massachusetts 02114, USA; GeneDx, Gaithersburg, Maryland, 20877, USA; Department of Pediatrics, Division of Genetics and Genomic Medicine, Washington University School of Medicine, St. Louis, MO, 63110, USA; Center for Statistical Genetics, Gertrude H. Sergievsky Center, Taub Institute for Alzheimer’s Disease and the Aging Brain, Department of Neurology, Columbia University Medical Center, 630 W 168th St, New York, NY 10032; University of Arkansas for Medical Sciences, Lowell, AK, 72745, USA; Department of Pediatrics, McKusick-Nathans Institute of Genetic Medicine, Johns Hopkins University, Baltimore, MD, 21205, USA

## Abstract

Germline pathogenic variants in chromatin-modifying enzymes are a common cause of pediatric developmental disorders. These enzymes catalyze reactions that regulate epigenetic inheritance via histone post-translational modifications and DNA methylation. Cytosine methylation of DNA (5mC) is the quintessential epigenetic mark, yet no human Mendelian disorder of DNA demethylation has been delineated. Here, we describe in detail the first Mendelian disorder caused by disruption of DNA demethylation. TET3 is a methylcytosine dioxygenase that initiates DNA demethylation during early zygote formation, embryogenesis, and neuronal differentiation and is intolerant to haploinsufficiency in mice and humans. Here we identify and characterize 11 cases of human *TET3* deficiency in 8 families with the common phenotypic features of intellectual disability/global developmental delay, hypotonia, autistic traits, movement disorders, growth abnormalities, and facial dysmorphism. Mono-allelic frameshift and nonsense variants in *TET3* occur throughout the coding region. Mono-allelic and bi-allelic missense variants localize to conserved residues with all but one occurring within the catalytic domain and most displaying hypomorphic function in a catalytic activity assay. *TET3* deficiency shows substantial phenotypic overlap with other Mendelian disorders of the epigenetic machinery, including intellectual disability and growth abnormalities, underscoring shared disease mechanisms.

## Introduction

Post-translational modifications of histone tails and DNA methylation play essential roles in development by regulating chromatin structure and gene expression. Inherited conditions that disrupt these processes – chromatin-modifying enzyme disorders or Mendelian disorders of the epigenetic machinery - account for a substantial percentage of neurodevelopmental and growth abnormalities in children ^1; 2^. Most known disorders in this class are caused by pathogenic variants in histone-modifying enzymes and chromatin remodelers. Far fewer have been linked to deficiencies in the DNA methylation machinery^3–5^. The latter include disorders caused by defects in DNA methylation “writers,” or DNA methyltransferases (DNMTs), such as immunodeficiency-centromeric instability-facial anomalies syndrome-1 (ICF syndrome) due to bi-allelic variants in *DNMT3B* (MIM: 242860), and Tatton-Brown-Rahman syndrome due to mono-allelic variants in *DNMT3A* (MIM: 615879), or by defects in reader proteins that bind to DNA methylation, such as Rett syndrome, which is caused by variants in *MECP2* (MIM: 312750) ^3–5^. No Mendelian disorder has been linked to the multi-step and tightly regulated process that removes DNA methylation.

The roles of DNMTs and proteins like MECP2 in “writing” and “reading” methyl marks on DNA have been known for decades, whereas the existence of enzymes that can actively reverse or “erase” DNA methylation was discovered more recently ^6; 7^. The ten-eleven translocase (TET) family of enzymes consists of three methylcytosine dioxygenases (TET1, TET2, and TET3) that initiate DNA demethylation through a series of sequential oxidation reactions converting 5-methyl cytosine (5mC) first to 5-hydroxymethylcytosine (5hmC) and then to 5-formylcytosine (5fC) and 5-carboxylcytosine (5caC), which are removed either passively by replication-dependent dilution or actively by thymidine DNA glycosylase ^8^. The process ultimately results in loss of the methylated base and replacement with an unmethylated cytosine ^6; 7; 9; 10^, effectively leading to DNA demethylation.

In addition to being an intermediate in the active removal of 5mC, 5hmC is suggested to have an independent role in gene regulation, though the exact nature remains unclear. Notably, 5hmC levels differ globally based on cell lineage and are particularly enriched in mammalian brains ^8; 11^. 5fC and 5caC are less well understood and may have unique functions as well^8^. *Tet3* is highly expressed in oocytes, zygotes, and neurons, and ablation of *Tet3* in mice leads to embryonic lethality ^8^. TET3 plays an important role in rapidly demethylating the paternal genome after fertilization, producing genome-wide increases in the oxidized 5mC intermediates 5hmC, 5fC, and 5caC ^12–16^. Importantly, *Tet3* haploinsufficiency causes neonatal sublethality or sub-Mendelian ratios in mice ^17^. Furthermore, inhibition or depletion of *Tet3* in mouse differentiated neurons can impact synaptic function ^18–20^. In humans, *TET3* is highly intolerant to loss-of-function alleles in control databases^21^, and homozygous missense variants in *TET3* were recently implicated as a possible cause for autosomal recessive intellectual disability in a single consanguineous family (further described here as family 3) ^22^. Together, these findings illustrate the important role of TET3 in early embryonic development and neuronal function.

Here, we provide the first detailed description of a cohort of individuals with a Mendelian disorder due to disruption of the DNA demethylation machinery, namely the TET3 enzyme. Whereas inheritance patterns vary and include both autosomal dominant and autosomal recessive forms, all affected individuals have in common a deficiency in TET3 function. This is either due to one or more missense variants within the highly conserved catalytic domain, most of which have been functionally validated to possess decreased TET3 activity, or to a single frameshift or nonsense variant. The phenotype is remarkably similar between affected individuals and is consistent with the broader group of Mendelian disorders of the epigenetic machinery, which often show global developmental delay/intellectual disability and other neurological manifestations, growth abnormalities, and characteristic craniofacial features ^1 23 24^.

## Results

Individual 1 presented with developmental delay, generalized overgrowth including macrocephaly, and some facial features reminiscent of Sotos syndrome (**Table 1**). Targeted testing for Sotos syndrome (MIM: 117550; *NSD1*) and the related Malan syndrome (MIM: 614753; *NFIX*), as well as methylation testing for Beckwith Wiedemann syndrome (MIM# 130650) and array comparative genomic hybridization (CGH), were negative. We performed research-based trio exome sequencing and identified bi-allelic rare variants in *TET3* (NM_001287491.1 c.2254C>T; p.Arg752Cys and c.3265C>A; p.Val1089Met). Through collaborations with other institutions and Genematcher ^25^ we subsequently identified an additional 10 affected individuals in 7 unrelated families with overlapping phenotypes and rare variants in *TET3* predicted to negatively impact catalytic function (**Table 1; Figure 1**).

**Figure 1.**
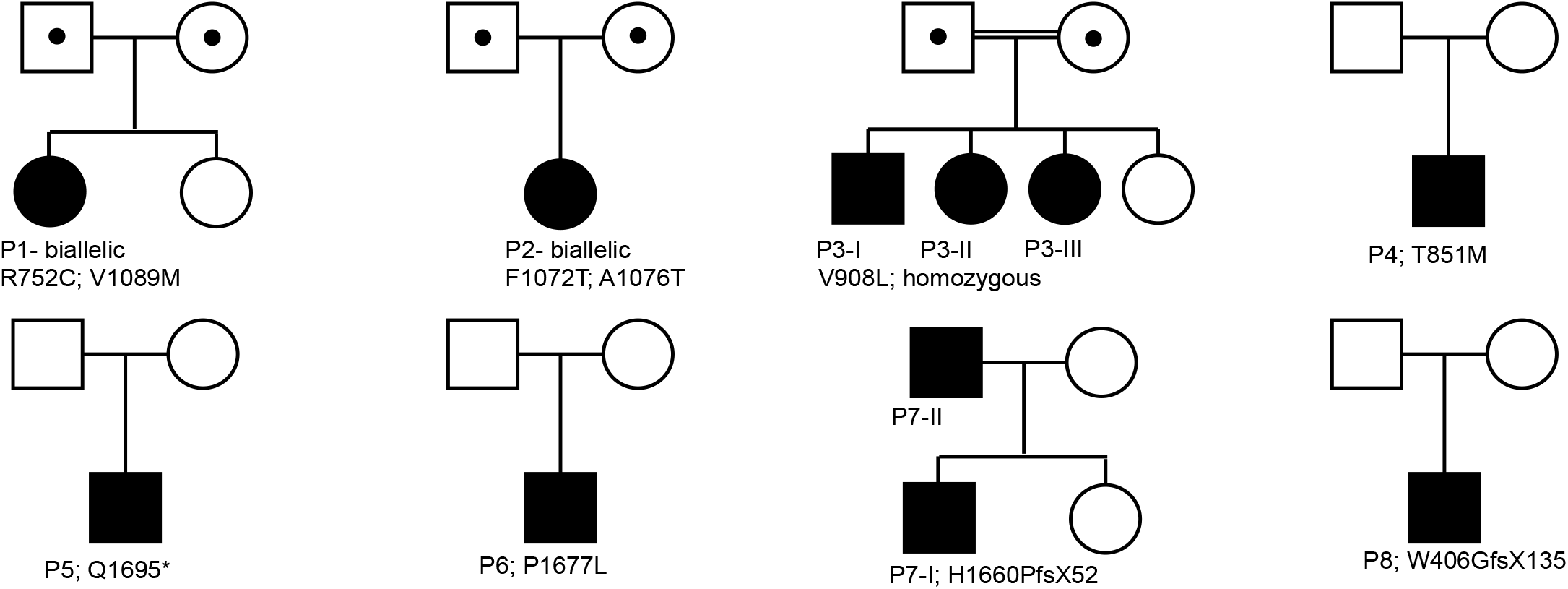
Inheritance patterns and variants in individuals with *TET3* deficiency. Pedigrees illustrating inheritance patterns in each family with specific variants listed.

**Table 1.**
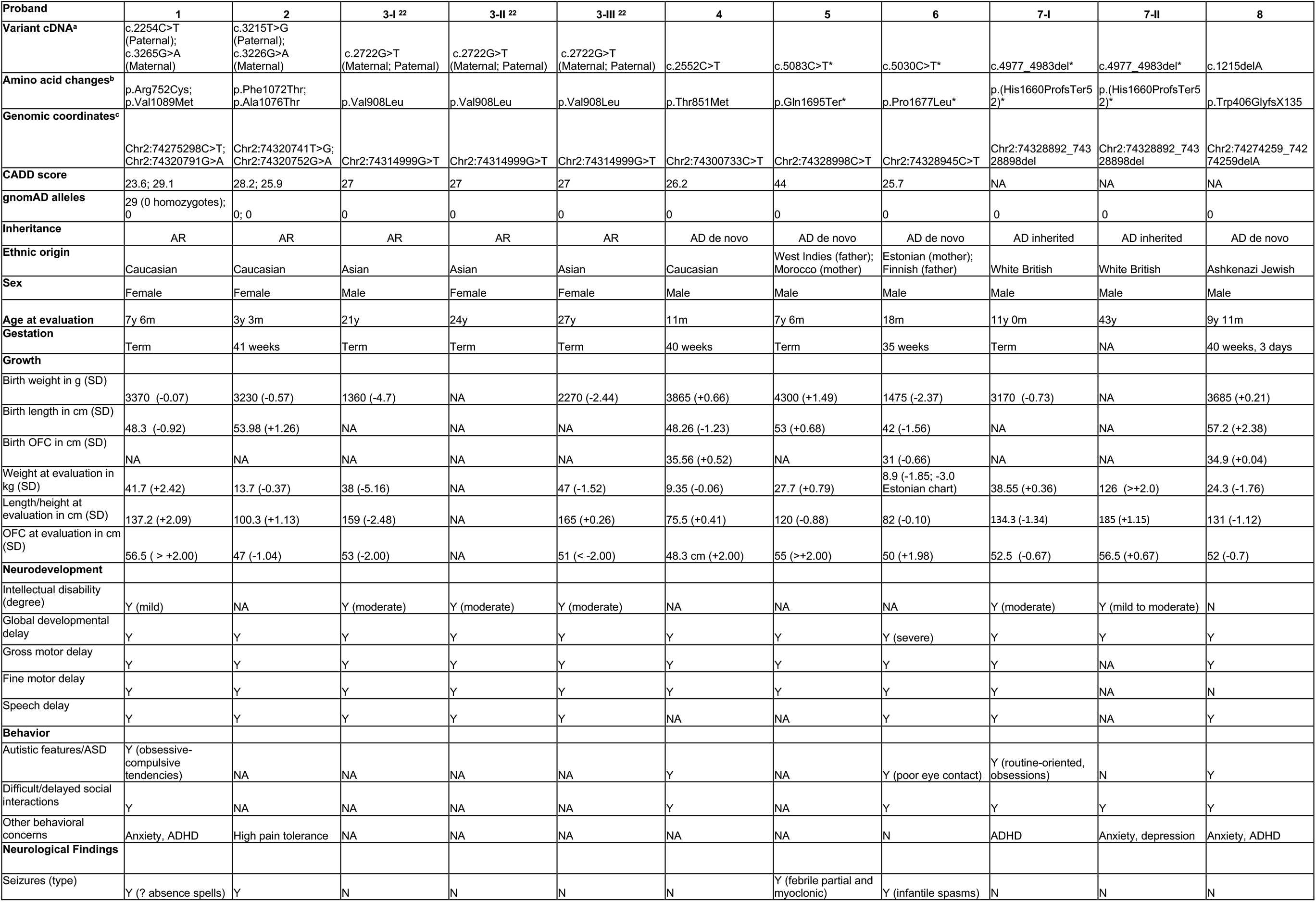
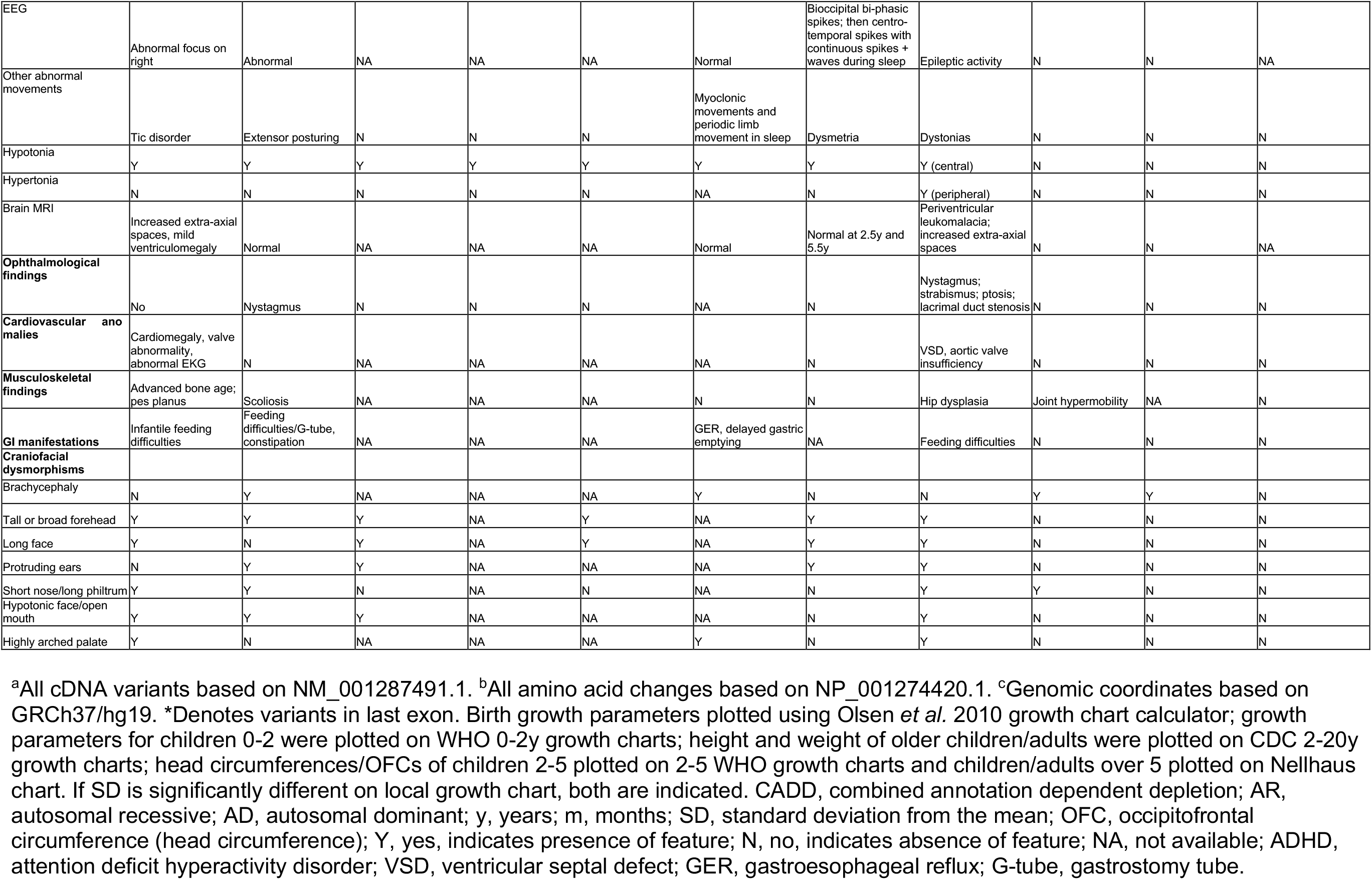
Clinical Characteristics of Individuals harboring *TET3* variants.

To delineate the phenotypic spectrum associated with variants in *TET3* we collected detailed clinical information on all affected individuals, who ranged in age from 11 months to 43 years at the time of assessment (**Table 1**). We observed striking phenotypic overlap among affected individuals (**Table 1**). All had global developmental delay and/or intellectual disability, and the vast majority had hypotonia and/or joint hypermobility (9/11). Other commonly observed findings were autistic features including difficulty with social interactions (6/11), growth abnormalities (8/11), movement disorders (5/11), and overlapping common facial characteristics **(Figure 1).** The developmental delay/intellectual disability ranged from mild to severe and included gross motor delay with or without speech delay in almost all cases (**Table 1**). Seizures and/or EEG abnormalities occurred in 4/11 individuals; other movement disorders were also noted and included tics, dystonia, extensor posturing, and myoclonic jerks (**Table 1**). Brain MRI demonstrated non-specific abnormalities in two individuals, including periventricular white matter changes and increased extra-axial spaces including ventriculomegaly (**Table 1**). Postnatal growth abnormalities were identified in 7/11 affected individuals, most often involving head size, with three individuals (1, 4, and 5) having true macrocephaly (OFC ≥ +2SD above the mean), one (6) having borderline/relative macrocephaly, and two (3-I and 3-III) having microcephaly (**Table 1**). In individual 1, macrocephaly is accompanied by tall stature (height ≥ +2SD above the mean), and in individual 3-I, microcephaly is accompanied by short stature (height ≤ −2SD below the mean; **Table 1**). Three individuals (3-I, 3-III, and 6) were born small for gestational age, suggesting a potential effect on prenatal growth; however, two of these were siblings from the same consanguineous family, and we cannot rule out other genetic causes, maternal factors, exposures, or poor prenatal care without additional information. The other individual born small for gestational age (individual 6) continued to exhibit poor weight gain but developed borderline macrocephaly by 18 months of age (**Table 1**). Distinctive craniofacial features common to these patients include tall and/or broad forehead (6/11) and long face (5/11; **Table 1**). Less commonly noted were brachycephaly (4/11), short nose and long philtrum particularly in younger individuals (4/11), hypotonic facies with open mouth appearance (4/11), protruding ears (4/11), and highly arched palate (3/11; **Table 1)**. A few had feeding difficulties (3/11), and eye findings including nystagmus (2/11; **Table 1**).

Five cases in three distinct families had bi-allelic variants, consistent with autosomal recessive inheritance (**Table 1; Figure 1**). Two of these individuals were compound heterozygotes, whereas the other three were siblings from a consanguineous family homozygous for the same variant (**Table 1; Figure 1**) ^22^. In all three of the autosomal recessive families, at least one parent appeared mildly affected. Both parents of individual 2 had mild learning difficulties requiring individualized educational plans (IEPs) in school; in addition, the father had attention deficit hyperactivity disorder (ADHD), and the mother had a history of seizures requiring medication in childhood. The mother of individuals 3-I, 3-II, and 3-III has severe anxiety, problems with short-term memory, and borderline psychosis, whereas the parental phenotypes in family 1 appear milder with the father and unaffected sister having specific and similar mild childhood learning disabilities, and the mother having occasional depression, significant anxiety, and possible ADHD, the latter two also confirmed in her affected daughter. All parents were able to live independently and/or hold jobs.

In terms of specific variants, all five autosomal recessive cases from three distinct lineages (**Figure 1**; **Table 1**) harbor either rare or novel missense changes at conserved residues within or adjacent to the catalytic domain of TET3 (**Figure 2A-B**), which consists of a dioxygenase domain separated by a spacer and a cystine-rich domain **(Figure 2A).** Specifically, individual 1 has a paternally inherited c.2254C>T (p.Arg752Cys) variant just upstream of the catalytic domain and a maternally inherited c.3265G>A (p.Val1089Met) variant within the dioxygenase domain; individual 2 has a paternally inherited c.3215T>G (p.Phe1072Thr) variant and a maternally inherited c.3226G>A (p.Ala1076Thr) variant, both of which are in the dioxygenase domain (**Figure 2A**). Individuals 3-I, 3-II, and 3-III are siblings from a consanguineous family and share the c.2722G>T (p.Val908Leu) homozygous variant^22^ within the cystine-rich domain (**Figure 2A**). We mapped these potential variants in *TET3* to the well-conserved TET2 catalytic domain crystal structure, and all but one (p.Arg752Cys) could be visualized **(Figure 2C)**. Val1089Met and Phe1072Thr are in close proximity to residue Asn1387 of TET2, which forms a hydrogen bond with the cytosine base of 5hmC to stabilize binding. Ala1076 is found adjacent to Thr1393 of TET2 which participates in hydrogen bonding with the N4 exocyclic amino group of cytosine ^26^.

**Figure 2.**
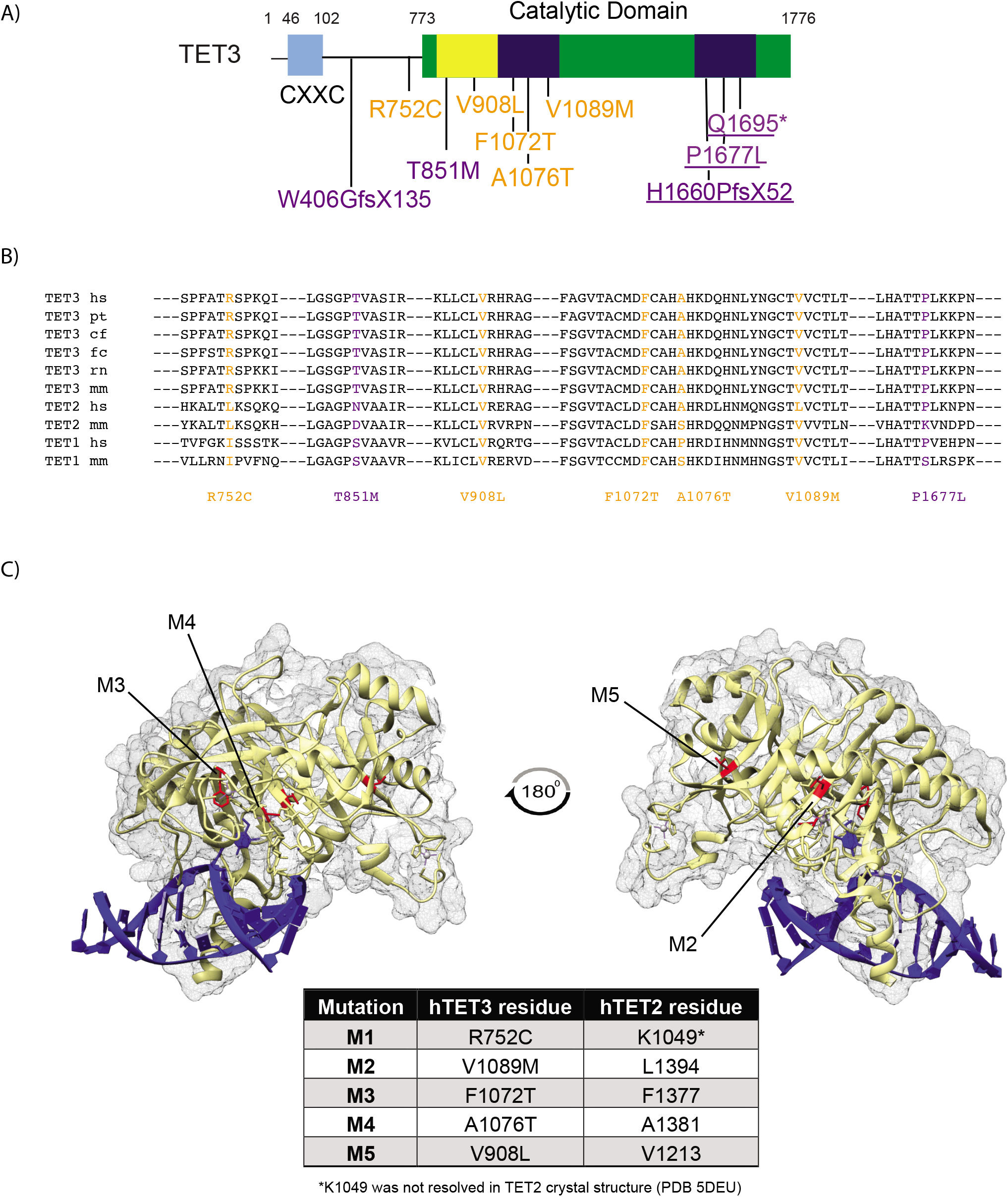
*TET3* variants and predicted functional consequences. **(A)** Schematic of TET3 protein showing domain structure with the catalytic dioxygenase domain in green (aa, amino acids 773-1776) and specific subdomains indicated as follows: the Cys-rich insert in yellow (aa 825-1012) and the double-stranded β helix domain in dark blue (DSBH; aa 1012-1159; aa 1636-1719). The DSBH domain is separated by a low complexity insert. The N-terminal CXXC DNA binding domain is shown in light blue (aa 46-102). Specific variants are annotated in orange for recessive alleles and purple for dominant alleles, and underlined variants occur within the last exon. **(B)** Alignment of missense variants in *TET3* across multiple species, including hs, *Homo sapiens;* pt, *Pan traglodytes;* cf, *Canis familiaris;* fc, *Felis catus;* rn, *Rattus norvegicus;* and mm, *Mus musculus*, and among TET enzymes. **(C)** Crystal structure of TET2 bound to DNA with highlighted *TET3* mutations M2, M3, M4, and M5.

In addition, we identified six individuals in five families with rare mono-allelic variants in *TET3* suggestive of autosomal dominant inheritance **(Table 1; Figure 1).** One was inherited, and the rest occurred *de novo*. In family 7, a similarly affected father and son both harbor the same frameshift variant c.4977_4983del (p.His1660ProfsTer52) in the catalytic domain (**Figure 2A**), consistent with autosomal dominant inheritance. Individual 5 has a *de novo* nonsense variant c.5083C>T (p.Gln1695Ter) also located within the dioxygenase domain, and both this and the inherited variant occur in the last exon (**Figure 2A**). Individual 8 has a *de novo* frameshift variant c.1215delA (p.Trp406GlyfsX135) upstream of the catalytic domain (**Figure 2A**), and individuals 4 and 6 harbor *de novo* missense variants, namely c.2552C>T (p.Thr851Met) and c.5030C>T (p.Pro1677Leu), with the former located in the cystine-rich domain and the latter within the dioxygenase domain (**Figure 2A**). Notably, in both autosomal recessive and autosomal dominant cases, all missense variants (except for p.Arg752Cys) are located within the catalytic domain (**Figure 2A**), and moreover, all occurred at residues highly conserved across species with many of the variants occurring at positions also conserved among human and sometimes mouse TET enzymes (**Figure 2B)**.

In the first step of DNA demethylation, 5mC is converted to 5hmC by TET enzymes ^8^. To analyze the effect of individual patient variants on TET3 catalytic activity, we measured 5hmC production using a cell culture system whereby recessively inherited TET3 variants (Arg752Cys, Phe1072Thr, Ala1076Thr, Val1089Met, and Val908Leu) were overexpressed in HEK 293 cells and total 5hmC levels were measured with a dot blot assay (**Figure 3A**). We compared the activity of TET3 variants to that of a known catalytically inactive mutant (Dcat; p.H1077Y/D1079A) and to full length wild-type TET3 **(Figure 3B-D)**. All patient variants tested demonstrated a defect in converting 5mC to 5hmC, except for Arg752Cys **(Figure 3 B,D)**, which is outside the catalytic domain and not conserved among TET enzymes **(Figure 2A-B)**. The observed defects were consistent across biological replicates, despite fluctuations in the levels of TET3 variant expression (**Figure S1**). For quantification, the 5hmC levels from cells expressing TET3 mutants were normalized to those measured in cells transfected with the wildtype TET3 construct to obtain a relative 5hmC signal **(Figure 3D)**. These results showing decreased cellular levels of 5hmC suggest that the vast majority of the missense variants identified in affected individuals have hypomorphic function. The observation that patients with nonsense and frameshift variants have phenotypes similar to patients with hypomorphic missense variants further supports the hypothesis that decreased TET3 catalytic activity leads to disease.

**Figure 3.**
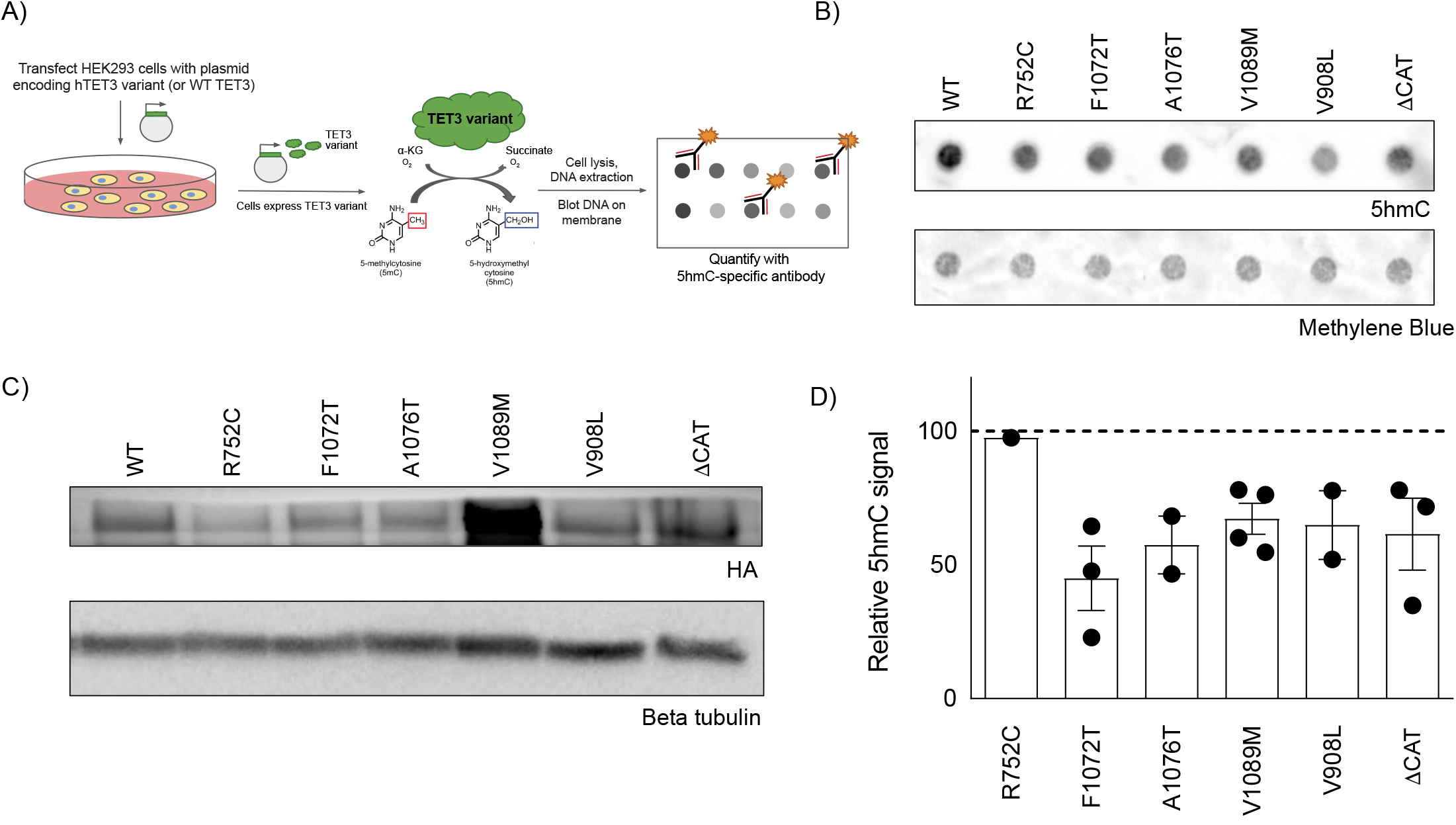
Cells overexpressing *TET3* variants have decreased levels of 5hmC. **(A)** Schematic outlining enzymatic activity assay for measuring TET3 catalytic activity. **(B)** Representative dot blot showing 5hmC levels in HEK293 cells overexpressing wild-type or mutant HA-tagged *TET3* constructs. **(C)** Representative Western blot showing wild-type and mutant TET3 protein expression in HEK293 cells. Tubulin was used as loading control. **(D)** Quantification of 5hmC signal relative to wild type. The dotted line indicates wild type signal. Error bars represent standard error of the mean. WT, wild type; Dcat, catalytically inactive control.

## Discussion

*TET3*, like most genes encoding components of the epigenetic machinery, is highly dosage-sensitive in both model organisms and humans^1; 17; 21^. *TET3* has a pLI score of 1 (observed/expected=0.02), suggesting near complete intolerance to loss of function variation ^21^. Based on this high pLI score and its high degree of coexpression across diverse tissues, TET3 was recently predicted bioinformatically to be a candidate epigenetic machinery disease gene^27^. Consistent with dosage sensitivity, most Mendelian disorders of the epigenetic machinery are autosomal dominant due to haploinsufficiency. In line with these observations, we identified patients with mono-allelic missense or loss of function (nonsense and frameshift) variants in *TET3*. However, we also report individuals with overlapping phenotypes that carry bi-allelic hypomorphic missense variants, each with mildly decreased catalytic activity according to our overexpression assay. We expect that in both cases - mono-allelic loss of function mutations and bi-allelic hypomorphic mutations - there is a similar reduction in total enzymatic activity causing a conserved disease mechanism across inheritance types. Notably, nonsense and frameshift variants were only identified in the heterozygous state, suggesting that some residual TET3 activity is required for viability. Conversely, perhaps a certain threshold of TET3 activity exists, below which developmental phenotypes result regardless of whether reduced TET3 activity is caused by missense or loss of function heterozygous alleles or biallelic hypomorphic alleles.

Within our cohort we identified patients with autosomal recessive inheritance with mildly affected carrier parents, which is consistent with a causal relationship between perturbations of TET3 activity and disease manifestations and suggests an inverse correlation between residual TET3 activity and phenotypic severity. Similar examples of complex inheritance involving both mono-allelic and bi-allelic variants in another component of the epigenetic machinery, *KDM5B*, have been reported recently ^28; 29^. An alternative hypothesis for differential modes of inheritance other than absolute activity levels could be that mono-allelic mutations have activating or dominant negative effects while bi-allelic mutations lead to loss of function; other examples of this exist in human disease ^30; 31^. We cannot rule this out here because the nonsense and frameshift variants identified in families 5 and 7, respectively, are located in the last exon and may escape non-sense mediated decay, raising the possibility of a dominant negative mutation mechanism and warranting further mechanistic studies to better correlate genotype and phenotype.

Along these lines, despite the evidence for multiple modes of inheritance, we cannot discount the possibility that individuals with *de novo* variants have additional non-coding sequence variants *in trans*, given that exome sequencing, not genome sequencing, was performed. Similarly, we have not ruled out epigenetic alterations *in trans*, such as DNA methylation. In addition, individuals can have additional variants that contribute to their phenotype as is the case for individual 8, with a paternally inherited 16p11.2 duplication and individual 2 with a maternally-inherited 16q22.1q22.2 duplication. Further studies, including genome sequencing and methylation analysis, could shed light on the molecular mechanisms involved. However, family 7 with two sequential affected generations supports autosomal dominant inheritance, particularly when considered along with the four *de novo* cases. Together, our observations strongly support two distinct modes of inheritance for *TET3* deficiency syndrome.

Another explanation for the observed monoallelic and biallelic cases is sex-specific differences. Notably, all probands with mono-allelic variants are male whereas all but one of the individuals with bi-allelic variants are female. While it remains true that most (if not all) of the carrier mothers appear to have mild manifestations, these were not sufficient to bring them to medical attention. It therefore remains possible that males are more susceptible to *TET3* deficiency and only require a single monoallelic variant to express the full phenotype whereas females require biallelic variants. Certainly, these sex-specific findings may be due to chance given the small total number of individuals; therefore, identification of additional affected individuals and further investigation into the mechanisms associated with specific mutations is required to fully delineate the mode of pathogenesis.

*TET3*-deficient individuals have significant phenotypic overlap with the broader group of Mendelian disorders of the epigenetic machinery, which are characterized by global developmental delay/intellectual disability and other neurobehavioral findings, as well as growth abnormalities including growth retardation or overgrowth ^1; 23; 24^. Tatton-Brown *et al*. recently showed that variants in epigenetic machinery genes account for approximately 45% of overgrowth co-occurring with intellectual disability and that the most common disorder within the group is Sotos syndrome, resulting from variants in *NSD1* ^23^. Our data suggest *TET3* deficiency may fall into this group of overgrowth and intellectual disability disorders, as all individuals with *TET3* deficiency have intellectual disability/global developmental delay and a subset have overgrowth. Among patients displaying both phenotypes, most also exhibit facial features reminiscent of Sotos syndrome, including a long face and tall forehead. This phenotypic overlap is particularly intriguing in the context of known biochemical interactions between TET3 and NSD proteins ^32^. Further support comes from observations that TET3 also binds other epigenetic factors encoded by genes responsible for overgrowth and intellectual disability disorders, such as *SETD2* responsible for Luscan-Lumisch syndrome (MIM: 616831), *HIST1H1E* responsible for Rahman syndrome (MIM: 617537), and *SUZ12*, which causes a newly described Weaver-like syndrome ^23; 32–34^. The subset of individuals with *TET3* deficiency who exhibit overgrowth (macrocephaly) in addition to intellectual disability is mostly confined to those with mono-allelic variants and autosomal dominant inheritance (individuals 4,5,6), consistent with the established inheritance pattern of overgrowth and intellectual disability disorders ^23^. The exception is individual 1, who has overgrowth consisting of macrocephaly and tall stature in the setting of bi-allelic variants inherited *in trans* from carrier parents, which we initially thought indicated autosomal recessive inheritance. However, her Arg752Cys variant was the only one tested that did not show decreased TET3 activity in our enzymatic assay, suggesting the possibility that the other p.Val1089Met variant is mostly if not solely responsible for the disease phenotype. In support of this, her mother, who carries the confirmed p.Val1089Met hypomorphic variant, does exhibit potentially relevant features, including significant anxiety and possible ADHD, although she is of average height and her head circumference measurement is unavailable. Further supporing this, the father and unaffected non-carrier sister have specific and similar mild childhood learning disabilities, which are likely unrelated to the proband’s phenotype. It therefore remains possible that the phenotype of individual 1 more closely resembles that of others with autosomal dominant inheritance. Future studies of individual mutation mechanisms will shed light on disease pathogenesis.

Together our clinical observations and biochemical studies define a novel neurodevelopmental disorder due to reduction in TET3 catalytic activity. TET3 plays a key role in actively reversing DNA methylation and is the first enzyme in the DNA demethylation system shown to cause a Mendelian disorder. Individuals with TET3 deficiency display overlapping phenotypic features with other Mendelian disorders of the epigenetic machinery, namely developmental delay/intellectual disability, other neurobehavioral manifestations, and growth abnormalities. By describing in detail for the first time a deficiency in the DNA demethylation pathway, our work defines a novel biochemical category of epigenetic machinery disorders and expands our knowledge of this important group diseases. Given the central role of DNA methylation in epigenetic inheritance, this disorder provides important initial insights into the dynamic regulation of DNA methylation in humans. Further characterization of *TET3*-deficient individuals, their causative variants, and their resulting molecular perturbations will lead to a deeper understanding of the role of DNA methylation and demethylation in human development and disease.

## Acknowledgements

We would like to thank all of the participating families. R.B. acknowledges support from the NIH (DP2MH107055, R01GM127408). A.P. was supported in part by an NIH training grant (T32 HD083185). J.A.F. acknowledges support from The Hartwell Foundation (Individual Biomedical Research Award) and the National Institutes of Health (K08HD086250). We thank the Baylor-Hopkins Center for Mendelian Genomics for exome sequencing and bioinformatics analysis on family 1; this work was supported by grant 5UM1HG006542 from NHGRI and NHLBI. Work on family 6 was supported by the Estonian Research Council grants PUT355, PRG471 and PUTJD827. The Broad Center for Mendelian Genomics (UM1 HG008900) is funded by the National Human Genome Research Institute with supplemental funding provided by the National Heart, Lung, and Blood Institute under the Trans-Omics for Precision Medicine (TOPMed) program and the National Eye Institute. Work on family 3 was supported by Higher Education Commission Grant No. NRPU-7099, Pakistan, to M.A. The DDD study presents independent research commissioned by the Health Innovation Challenge Fund [grant number HICF-1009-003]. This study makes use of DECIPHER (http://decipher.sanger.ac.uk), which is funded by the Wellcome. See Nature PMID: 25533962 or www.ddduk.org/access.html for full acknowledgement.

## Methods

### Human subjects

Written informed consent was obtained from all individuals or family member legal representatives prior to exome sequencing. Individual 1 was counseled regarding the possible outcomes of exome sequencing and signed a consent form for research-based exome sequencing through the Baylor-Hopkins Center for Mendelian Genomics, which was approved by the Johns Hopkins Institutional Review Board (IRB). The rest of the participants were recruited through GeneMatcher ^25^. Individuals 2 and 4 were consented for clinical exome sequencing through Greenwood Genetic Center, and individual 8 was consented for clinical exome sequencing through GeneDx. Individuals 5 and 6 were consented for clinical and/or research-based exome sequencing. Individuals 3-I, 3-II, and 3-III were consented for research-based exome sequencing as described^22^, and individuals 7-I and 7-II were consented for research-based trio exome sequencing through the Deciphering Developmental Disorders (DDD) study^35^.

### Exome and Sanger sequencing

For individual 1, trio exome sequencing was performed on genomic DNA isolated from saliva through the Baylor-Hopkins Center for Mendelian Genomics at Johns Hopkins. Bi-allelic rare variants in *TET3* were identified using standard bioinformatics analysis. Sanger sequencing confirmed the presence of the *TET3* variants in the trio and their absence in an unaffected sibling. Individuals 2 and 4 had trio exome sequencing performed at Greenwood Genetic Center on a clinical basis. Standard bioinformatics analysis revealed bi-allelic (individual 2) and mono-allelic (individual 4) rare variants in *TET3*, which were subsequently confirmed by Sanger sequencing. Individuals 3-I, 3-II, and 3-III had exome sequencing performed as described^22^. Individual 5 had trio exome sequencing performed with standard bioinformatics analysis, which identified a *de novo* monoallelic variant in *TET3*; the variant was confirmed with Sanger sequencing. Individual 6 had clinical trio exome sequencing performed, which did not reveal pathogenic mutations in genes known to be associated with Mendelian disorders. Reanalysis at the Broad Institute of MIT and Harvard identified a rare *de novo TET3* missense variant, which was subsequently confirmed with Sanger sequencing. Individual 7 had trio exome sequencing performed as part of the DDD study ^35^; the rare inherited mono-allelic variant in *TET3* was identified in the proband and his similarly-affected father using standard bioinformatics analysis. Sanger sequencing confirmed the presence of the *TET3* variant in the affected proband and his father and its absence in the mother and an unaffected sibling. Individual 8 had trio exome sequencing performed on a clinical basis through GeneDx. Standard bioinformatics analysis was performed and revealed a *de novo* mono-allelic variant in *TET3*, which was confirmed by Sanger sequencing. All patients reported have no known definitive pathogenic variants identified in genes causative for developmental delay.

### Cells

HEK 293 cells were cultured in Dulbecco’s modified Eagle’s medium (DMEM; Gibco) containing 10% fetal bovine serum (FBS; Sigma), 2 mM L-Glutamine (Gibco), MEM Non-essential amino acid solution (Sigma), sodium pyruvate (Sigma), and Penicillin Streptomycin.

### Cloning and plasmids

Full-length human TET3 coding sequence was amplified from cDNA and cloned into the pINTO-N3 plasmid backbone. The pINTO-N3 vector was based on the pINTO system ^36^, containing three N-terminal epitope tags (FLAG, HA, and Twin-Strep-Tag). Point mutations were introduced into the *hTET3* coding sequence via Gibson assembly and verified by Sanger sequencing.

### 5hmC dot blot

HEK 293 cells were transiently transfected with plasmids using Lipofectamine 3000 (Thermo Fisher). Cells were harvested 48 hours post transfection, lysed in Proteinase K buffer (100 mM NaCl, 10 mM Tris-HCl pH 8.0, 10mM EDTA, 0.5% SDS), and sonicated for 20 minutes (30 seconds ON, 30 seconds OFF) using a Bioruptor (Diagenode, NJ). A portion of the cell lysate was used to perform a Western blot with an HA antibody (#901501 BioLegend, CA) to verify hTET3 protein expression. The remaining cell lysate was incubated with Proteinase K (Ambion AM2548) for 45 minutes at 50°C and DNA was extracted using Phenol-Chloroform Isoamyl Alcohol (PCIA). DNA was denatured for 10 minutes at 95°C in 100 mM NaOH and 10 mM EDTA and then neutralized by addition of 2M ammonium acetate (pH 7.0). Denatured DNA was transferred to a BrightStar-Plus Nylon membrane (Ambion AM10102) using a Bio-Dot microfiltration apparatus (Bio-Rad) according to the manufacturer’s instructions. Briefly, the membrane was rinsed in 6X SSC buffer before assembling the apparatus and the DNA samples were loaded onto the membrane under vacuum pressure. The membrane was rinsed in 2X SSC and then dried for 15 minutes at 80°C in a hybridization oven. The membrane was then crosslinked twice with 200 mJ/cm^2^ UVA (254 nm) using a Spectrolinker (Spectroline, NY). Next, the membrane was stained with a 0.04% solution of Methylene Blue (Sigma 66720) to visualize total DNA. A Western blot was then performed to detect 5hmC using a 5hmC antibody (Active Motif 39770).

### Dot blot quantification and analysis

Dot blot quantification was performed using ImageJ as described in the ImageJ documentation. Raw TIFF files were opened in ImageJ and the “Integrated Density” measurement of each dot was recorded after correcting for background. To account for potential differences in total DNA amount across samples, the 5hmC signal was divided by the Methylene Blue signal for each dot. The normalized 5hmC signal was averaged across biological replicates and divided by the normalized 5hmC signal in the wild-type *hTET3* transfection to obtain a relative 5hmC signal. Samples in which the transfected mutant *hTET3* was expressed at lower levels than the wild-type control were not further considered in the analysis. These were the only data points excluded from the final quantification.

### Mutation Modeling

Mutations in *TET3* were mapped onto the well-conserved TET2 catalytic domain crystal structure (PDB accession 5DEU ^26^) using UCSF Chimera ^37^.

## Web Resources

CADD, http://cadd.gs.washington.edu/

ClinVar, https://www.ncbi.nlm.nih.gov/clinvar/

gnomAD Browser, http://gnomad.broadinstitute.org/

OMIM, http://www.omim.org/

PyMOL, https://pymol.org/2

RCSB Protein Data Bank, http://www.rcsb.org/pdb/home/home.do

