## Supplemental Figures for "Delineation of the First Human Mendelian Disorder of the DNA Demethylation Machinery: *TET3* Deficiency"

**Figure S1**

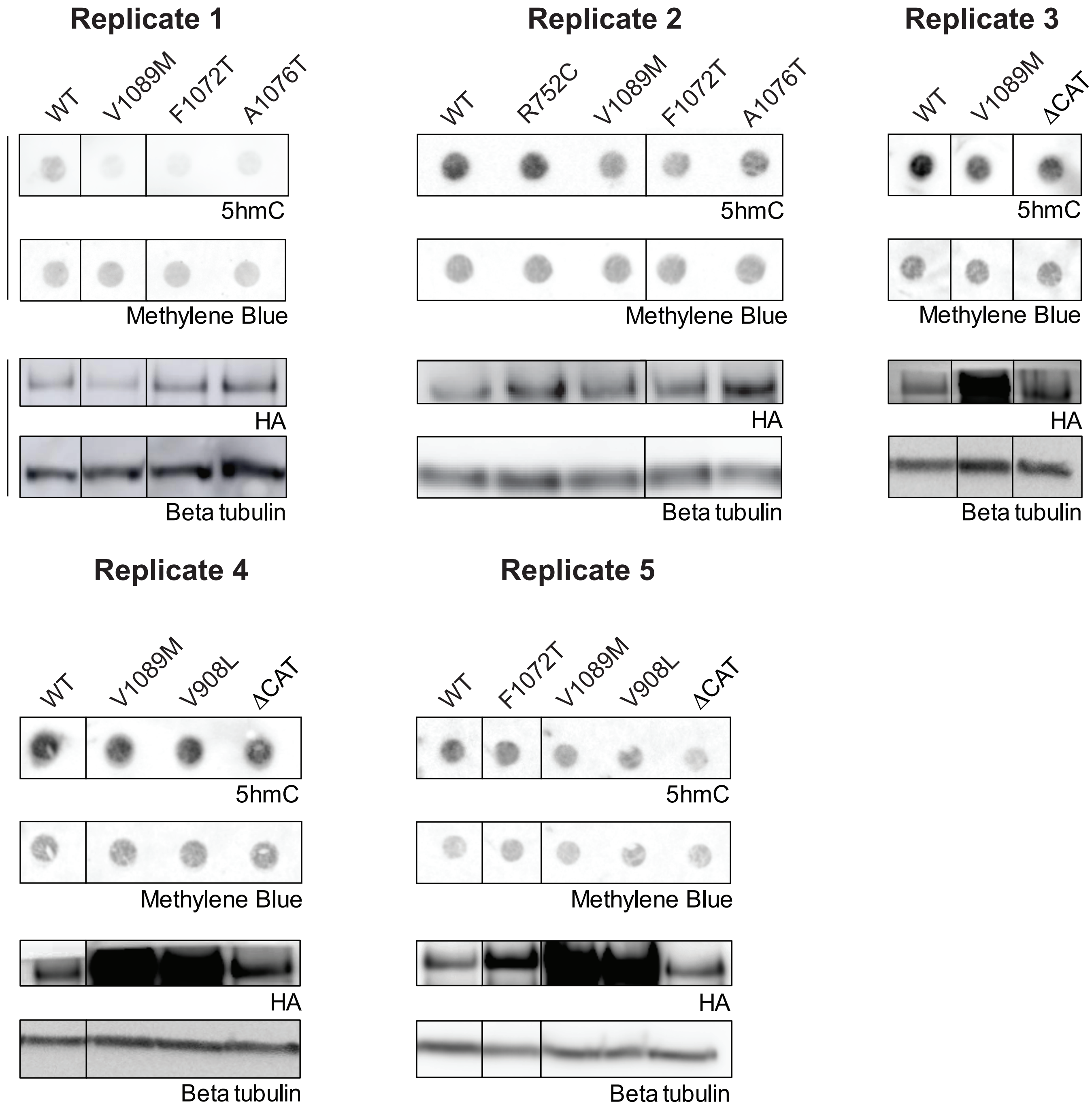


**Figure S1. 5hmC dot blots and Western blots used for final analysis.** 5hmC dot blot and Western blot images from five biological replicates. Samples that were not used for the final quantification in Figure 3 are not displayed.
